# A Kirchhoff-Nernst-Planck framework for modeling large scale extracellular electrodiffusion surrounding morphologically detailed neurons

**DOI:** 10.1101/261107

**Authors:** Andreas Solbrå, Aslak Wigdahl Bergersen, Jonas van den Brink, Anders Malthe-Sørenssen, Gaute T. Einevoll, Geir Halnes

## Abstract

1

Many pathological conditions, such as seizures, stroke, and spreading depression, are associated with substantial changes in ion concentrations in the extracellular space (ECS) of the brain. An understanding of the mechanisms that govern ECS concentration dynamics may be a prerequisite for understanding such pathologies. To estimate the transport of ions due to electrodiffusive effects, one must keep track of both the ion concentrations and the electric potential simultaneously in the relevant regions of the brain. Although this is currently unfeasible experimentally, it is in principle achievable with computational models based on biophysical principles and constraints. Previous computational models of extracellular ion-concentration dynamics have required extensive computing power, and therefore have been limited to either phenomena on very small spatiotemporal scales (micrometers and milliseconds), or simplified and idealized 1-dimensional (1-D) transport processes on a larger scale. Here, we present the 3-D Kirchhoff-Nernst-Planck (KNP) framework, tailored to explore electrodiffusive effects on large spatiotemporal scales. By assuming electroneutrality, the KNP-framework circumvents charge-relaxation processes on the spatiotemporal scales of nanometers and nanoseconds, and makes it feasible to run simulations on the spatiotemporal scales of millimeters and seconds on a standard desktop computer. In the present work, we use the 3-D KNP framework to simulate the dynamics of ion concentrations and the electrical potential surrounding a morphologically detailed pyramidal cell. In addition to elucidating the single neuron contribution to electrodiffusive effects in the ECS, the simulation demonstrates the efficiency of the 3-D KNP framework. We envision that future applications of the framework to more complex and biologically realistic systems will be useful in exploring pathological conditions associated with large concentration variations in the ECS.

**Author summary:** Many pathological conditions, such as epilepsy and cortical spreading depression, are linked to abnormal extracellular ion concentrations in the brain. Understanding the underlying principles of such conditions may prove important in developing treatments for these illnesses, which incur societal costs of tens of billions annually. In order to investigate the role of ion-concentration dynamics in the pathological conditions, one must measure the spatial distribution of all ion concentrations over time. This remains challenging experimentally, which makes computational modeling an attractive tool. We have previously introduced the Kirchhoff-Nernst-Planck framework, an efficient framework for modeling electrodiffusion. In this study, we introduce a 3-dimensional version of this framework and use it to model the electrodiffusion of ions surrounding a morphologically detailed neuron. The simulation covered a 1 mm^3^ cylinder of tissue for over a minute and was performed in less than a day on a standard desktop computer, demonstrating the framework’s efficiency. We believe this to be an important step on the way to understanding phenomena involving ion concentration shifts at the tissue level.

## 3 Introduction

The brain mainly consists of a dense packing of neurons and neuroglia, submerged in the cerebrospinal fluid which fills the *extracellular space* (ECS). Neurons generate their electrical signals by exchanging ions with the ECS through ion-selective channels in their plasma membranes. During normal signaling, this does not lead to significant changes in local ion concentrations, as neuronal and glial transport mechanisms work towards maintaining ion concentrations close to baseline levels. However, endured periods of enhanced neuronal activity or aberrant ion transport may lead to changes in ECS ion concentrations. Local concentration changes often coincide with slow shifts in the ECS potential [1–3], which may be partly evoked by diffusive electrical currents, i.e., currents carried by charged ions moving along ECS concentration gradients [2, 4]. While concentration gradients can influence electrical fields, the reverse is also true, since ions move not only by diffusion but also by electric drift. A better understanding of the electrodiffusive interplay between ECS ion dynamics and ECS potentials may be a prerequisite for understanding the mechanisms behind many pathological conditions linked to substantial concentration shifts in the ECS, such as epilepsy and spreading depression [3, 5–7].

A simultaneous and accurate knowledge of the concentration of all ion species is needed to make reliable estimates of electrodiffusive effects in the ECS. Although this is currently unfeasible experimentally, it is in principle achievable with computational models based on biophysical principles and constraints. However, in most computational models in neuroscience ion-concentration dynamics are only partially modeled, or are ignored altogether. One reason for this is the challenge involved in keeping track of all ion concentrations and their spatiotemporal dynamics. Another reason may be the strong focus within the community on modeling the neuronal membrane dynamics at short timescales, during which both intra- and extracellular concentration changes are relatively small and putatively negligible. For example, the most common computational models for excitable cells, the multi-compartmental models and the cable equation, are based on the assumptions that (i) the ECS potential is constant (ground), and (ii) the ion concentrations are constant [8, 9]. The NEURON simulator [10, 11] is based on these assumptions, and although they are physically incorrect, they still allow for efficient and fairly accurate predictions of the membrane-potential dynamics.

Because of assumption (i), multi-compartmental models are unsuited for modeling ECS dynamics, and several approaches have been taken to construct models which include ECS effects. A majority of computational studies of ECS potentials are based on *volume conductor* (VC) theory [12–17]. VC-schemes link neuronal membrane dynamics to its signatures in the ECS potential. In contrast to the multi-compartmental models, VC-schemes are derived by allowing the ECS potential to vary, but still assuming that the ion concentrations are constant. VC-schemes are attractive, because they offer closed-form solutions, and allow the calculation of the electric field for arbitrarily large systems. Although it may be reasonable to neglect variations in ECS ion concentrations on short timescales, the accumulative effects of endured neuronal activity may lead to significant concentration changes in the ECS, which are related to the aforementioned pathological conditions. Naturally, models that do not include ion-concentration dynamics are not applicable for exploring such pathologies. Furthermore, VC-schemes neglect the effects from diffusive currents on the ECS potentials [4, 18, 19], and in previous computational studies we have found the low-frequency components of the ECS potential to be dominated by diffusion effects [4, 20].

VC-schemes include electric effects in a simplified manner, but ignore ion-concentration dynamics completely. A simplified approach to modeling concentration dynamics in brain tissue, is to use reaction-diffusion schemes (see e.g., [21–23]). In these schemes, concentration dynamics are simulated under the simplifying assumption that ions move due to diffusion only. This approach has been used for many specific applications, giving results in close agreement with experimental data [22]. However, the net transport of abundant charge carriers such as Na^+^, K^+^, Ca^2+^, and Cl^−^, is also influenced by electric forces, which is not incorporated in diffusion only (DO)-schemes. Furthermore, DO-schemes do not include the influence that diffusing ions can have on the electrical potential.

To account for the electric interactions between the different ion species, as well as the effect of such electric forces on the ECS potential, an electrodiffusive modeling framework is needed. The most detailed modeling scheme for electrodiffusion is the Poisson-Nernst-Planck (PNP) scheme [24–30]. The PNP-scheme explicitly models charge-relaxation processes, that is, tiny deviations from electroneutrality involving only about 10^−^9 of the total ionic concentration [31]. This requires a prohibitively high spatiotemporal resolution, which makes the PNP-scheme too computationally expensive for modeling the ECS on the tissue scale. Even the state-of-the-art simulations in the literature are on the order of milliseconds on computational domains of micrometers. The PNP-scheme is therefore not suited for simulating processes taking place at the tissue scale [19].

A series of modeling schemes have been developed that circumvent the brief charge-relaxation processes, and solve directly for the ECS potential when the system is in a quasi-steady state [4, 19, 32–38]. Circumventing charge-relaxation allows for simulations on spatiotemporal scales which are larger, compared to what is possible with the PNP-scheme, by several orders of magnitude. The charge-relaxation can be bypassed by replacing Poisson’s equation with the constraint that the bulk solution is electroneutral. These schemes have been shown to deviate from the PNP-scheme very close to the cell membrane (less than 5 µm), but to give a good agreement in the bulk solution [19]. The simplest electroneutral modeling scheme is the *Kirchhoff-Nernst-Planck (KNP)* scheme, previously developed in our group [37, 38]. A similar scheme was developed in parallel in the heart cell community [36].

The KNP-scheme has previously been used to study electrodiffusive phenomena such as spatial K^+^ buffering by astrocytes [37], effects of ECS diffusion on the local field potential [4], and the implication for current-source density analysis [20]. For simplicity, these previous applications were limited to idealized 1-D setups with a relatively coarse spatial resolution. Furthermore, a comparison between the KNP framework and other simulation frameworks was not included in previous studies.

In the present study, we introduce a 3-D version of the KNP framework which can be used to simulate the electrodiffusive dynamics of ion-concentrations and the electrical potential in the ECS surrounding a morphologically detailed neuron, on large spatiotemporal scales. We further establish in which situations the KNP-scheme is accurate by comparing it to the more physically detailed PNP-scheme, and identify the conditions under which an electrodiffusive formalism is needed by comparing the KNP-scheme to the less detailed VC- and DO-schemes.

We present the results of three distinct simulation setups, which we will refer to as Application 1, Application 2, and Application 3 for the remainder of this study. The first two applications are simplified simulation setups, used to better understand the differences between the schemes introduced above:

1. An idealized 1-D domain filled with a salt solution, starting with a nonzero ion concentration gradient. We solve the system using the PNP-scheme, the KNP-scheme, and a DO-scheme. We compare results on short and long timescales (nanoseconds and seconds), to highlight the similarities and differences between the schemes.
2. A 3-D domain with an ion concentration point source and a point sink, of equal magnitude, embedded in a standard ECS ion solution. We compare results obtained with the VC- and KNP-schemes to highlight their similarities and differences.

The final application, which is the main result of this study, illustrates the scales at which the KNP-scheme can be used:

3. A morphologically detailed neuron embedded in a 3-D ECS solution. The ECS dynamics is computed using the KNP-scheme, and show how concentration gradients gradually built up in the space surrounding the active neuron, and how this influenced the local potential in the ECS. We compare results obtained with the VC-, DO-, and KNP-schemes to highlight their similarities and differences.

To our knowledge, the KNP-scheme is the first simulation framework which can handle 3-D electrodiffusion in neuronal tissue at relatively large spatiotemporal scales without demanding an insurmountable amount of computer power. For Application 3, the long-term ECS ion-concentration dynamics (about 100 s) in a spatial region of about 1 mm^3^ was run on a standard desktop computer within a day. We expect that the presented simulation framework will be of great use for future studies, especially for modeling tissue dynamics in the context of exploring pathological conditions associated with large shifts in ECS ion concentrations [3, 5–7].

## 4 Materials and methods

This section is thematically split into three parts. We begin by explaining the necessary physical theory, stating and deriving the equations which we implemented. Then, we explain in more detail how the models were implemented, including details such as numerical schemes and boundary conditions. Finally, we give the specific details on each of the three applications used in the study. The source code can be found online, at https://github.com/CINPLA/KNPsim, and the results in this study can be reproduced by checking out the tag PLoS.

### 4.1 Theory

#### 4.1.1 The Nernst-Planck equation for electrodiffusion

The ion concentration dynamics of an ion species in a solution is described by the continuity equation:

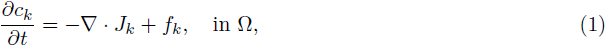

where *c_k_* is the concentration of ion species *k*, *f_k_* represent any source terms in the system, Ω is the domain for which the concentrations are defined, and *J_k_* is the concentration flux of ion species *k*. In the Nernst-Planck equation, *J_k_* consists of a diffusive and an electric component:

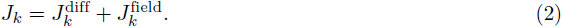

The diffusive component is given by Fick’s first law,

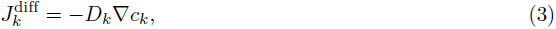

where *D_k_* is the diffusion coefficient of ion species *k*. The electric component is

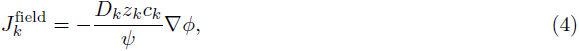

where *ϕ* is the electric potential, *z_k_* is the valency of ion species *k*, and *ψ* = *RT/F* is a physical constant, where *R* is the gas constant, *T* is the temperature, and *F* is Faraday’s constant (cf. Tab 1). Inserting Eqs 2-4 into Eq 1, yields the time evolution of the concentration of ion species *k*:

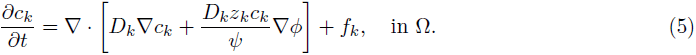

**Table 1.** The physical parameters used in the simulations.

| symbol | explanation | value |
| --- | --- | --- |
| $R$ | gas constant | 8.314 J/(K mol) |
| $T$ | temperature | 300 K |
| $F$ | Faraday's constant | $9.648 \times 10^4$ C/mol |
| $\epsilon_0$ | vacuum permittivity | $8.854 \times 10^{-12}$ F/m |
| $\epsilon_r$ | relative permittivity | 80 |

We model the ECS as a continuous medium, while in reality, the ECS only takes up roughly 20 % of the tissue volume [40] in the brain. To compensate for this, we use the porous medium approximation [41]. This involves two changes to the model. The diffusion constants of the ion species are modified as

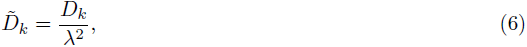

where *λ* is the tortuosity of the ECS. We used the value *λ* = 1.6 [42]. We denote the fraction of tissue volume belonging to the ECS by *α*, and set the value *α* = 0.2. The sources in the system are modified as

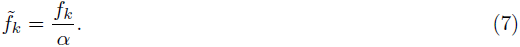

*D∼_k_* and *f∼_k_* are the values used in the simulations, and we will refer to these in the remainder of this study. In this study, we included four ion species: Ca^2+^, K^+^, Na^+^, and a general anion X^−^ accounting for all negative ions in the system. Their diffusion constants, as well as their steady-state values assumed for the ECS, are shown in Table 2.

**Table 2.** Diffusion constants and baseline ECS concentrations for the ion species considered. All ion constants were modified as *D∼_k_* = *D_k_/λ*^2^, where *λ* = 1.6 is the tortuosity. The general anion X^−^ was given the properties of Cl^−^.

|  | Na<sup>+</sup> | K<sup>+</sup> | X<sup>-</sup> | Ca<sup>2+</sup> |
| --- | --- | --- | --- | --- |
| $D$ ($\mu\text{m}^2/\text{ms}$) | 1.33 | 1.96 | 2.03 | 0.71 |
| $c_{\text{out}}$ (mM) | 150 | 3 | 155.8 | 1.4 |

#### 4.1.2 Poisson-Nernst-Planck (PNP) framework

In order to solve the Nernst-Planck equation, we need an expression for the electrical potential. One common approach is to assume Poisson’s equation for electrostatics,

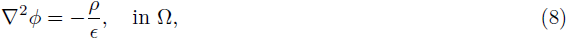

where *ρ* is the charge concentration in the system, and *ε* is the permittivity of the system, given by *ε* = *ε*_*r*_*ε*_0_, where *ε*_0_ is the vacuum permittivity, and *ε*_*r*_ is the relative permittivity of the extracellular medium (cf. Table 1).

The charge concentration is given by the sum of contributions from the different ion species,

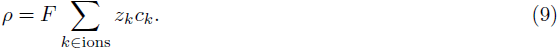

Poisson’s equation (Eq 8) combined with the Nernst-Planck equation (Eq 5) is referred to as the Poisson-Nernst-Planck equations. The PNP-system is defined at all points in space and gives physically correct results in cases where the continuum approximation for the ions is valid. However, there are a few challenges involved in solving the PNP system. Firstly, when neuronal membranes are present in the system, these need to be defined in terms of appropriate boundary conditions. Secondly, the PNP system is numerically very inefficient, because it models charge-relaxation processes which in the tissue solution take place at the spatiotemporal scales of nanometers and nanoseconds [43, 44].

#### 4.1.3 Kirchhoff-Nernst-Planck (KNP) framework

Several frameworks that assume the system to be electroneutral at all interior points have been developed to overcome the limitations of the PNP framework [19, 32{37]. We will here present one of these frameworks, which we have coined the Kirchhoff-Nernst-Planck framework.

In the KNP framework, the electric field is required to be such that:

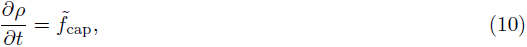

at all points in the system. In KNP, *f∼*_cap_ is a source term that exclusively stems from the capacitive current across a neuronal membrane, and which is zero anywhere else. Hence, the bulk solution is assumed to be electroneutral, so that *∂ρ/∂t* = 0. In other words, the only allowed nonzero charge density in the KNP-system is that building up the membrane potential across a capacitive membrane.

The rationale behind assuming bulk electroneutrality is that (i) charge concentration will only deviate from zero on nanometer scales, and that (ii) charge-relaxation occurs within a few nanoseconds, after which the system will settle at a quasi-steady state where *∂ρ/∂t* ≈ 0. Furthermore, the number of ions that constitute the net charge density at equilibrium is about nine orders of magnitudes smaller than the number of ions present [31]. Hence, if we simulate neuronal processes that take place at the spatiotemporal resolution of, or larger than, micrometers and microseconds, we would expect the bulk solution to appear electroneutral.

Motivated by this, the KNP-scheme bypasses the rapid equilibration process by assuming that the quasi-steady state is reached instantaneously, and derives the value for *ϕ* associated with the equilibrium state. In doing so, the KNP-scheme implicitly neglects the tiny local charge separation associated with the charge-relaxation process. We investigate the magnitude of this charge separation in the Results-section.

To turn Eq 10 into an equation which can be solved for *ϕ*, we insert the Nernst-Planck equation. Inserting Eq 9 into Eq 10 gives

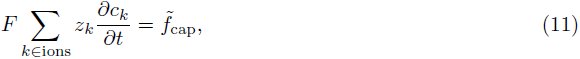

and inserting Eq 5 gives

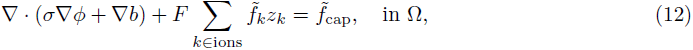

where *σ* is the conductivity of the medium, defined as [45]:

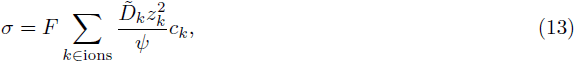

and

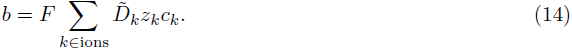

Eq 12 is similar to Poisson’s equation, in that it is an elliptic equation that can be solved for *ϕ*, assuming that *c_k_* is known.

The potential can be separated into the contribution from diffusive effects, and the contribution from the membrane currents. In order to analyze these components separately, we replace Eq 12 with the following equivalent set of equations:

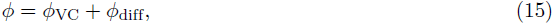

where

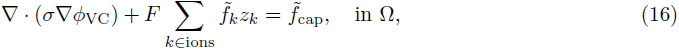

and

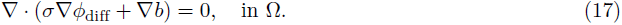

#### 4.1.4 Volume Conductor (VC) framework

Standard VC-schemes include only the effects of the transmembrane currents, and ignore all other ion-concentration dynamics. In VC-schemes, the electric potential is found by Eq 16. As the ECS concentrations are not modeled, *σ* is typically defined to be constant in VC-schemes, and current sources are modeled as either point or line sources [46]. This approach gives a closed-form solution for *ϕ*, which means it can be applied to arbitrarily large systems. In this study, when we refer to *ϕ*_VC_, we mean *ϕ*_VC_ as found in the KNP-scheme (Eq 16). This is essentially the same as in most common VC-implementations, with the exception that *σ* in Eq 16 is found from the ion concentrations.

#### 4.1.5 Diffusion Only (DO) framework

Reaction-diffusion schemes ignore all electric forces in the system. In our implementation, we obtained this by setting *ϕ* = 0 in Eq 5. The resulting equation for the concentration dynamics is:

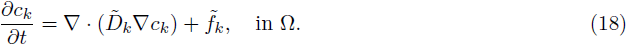

This is the equation used when we refer to the DO-scheme in the Results-section.

### 4.2 Implementation

The solver for the above modeling schemes was implemented utilizing FEniCS, an open-source platform for solving partial differential equations using the finite element method [39]. The reader is referred to [47] for an extensive introduction to the finite element method. The time derivative was approximated using an implicit Euler time-stepping scheme. We chose to use this method as the PNP equations are highly unstable, and the implicit Euler step offers superior stability to other methods [29]. Employing the implicit Euler step makes the PNP-scheme fairly stable, at the cost of numerical efficiency. Piecewise linear Lagrangian elements were used for all unknowns in all simulations [47].

The system was solved monolithically, using Newton’s method to solve for each time step. Due to limitations of FEniCS’ built-in nonlinear solver, we implemented our own Newton solver, which is found in the source code.

#### 4.2.1 Boundary conditions

For the first application, we employed a sealed (no-flow) boundary for the ion concentrations,

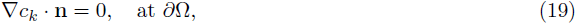

where **n** is a unit vector directed out of the domain, and *∂*Ω is the boundary of the domain.

For the second and third application, we assumed a concentration-clamp boundary condition,

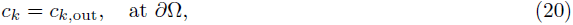

where *c_k,_*_out_ was set to typical ECS baseline concentrations, see Table 2. This can be interpreted as our system interacting with a larger reservoir of ions (the rest of the brain).

In order to maintain overall electroneutrality in the system, the KNP-scheme dictates that no net charge can leave the system on the boundary. To ensure this, we require that

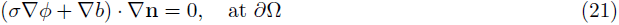

In order to compare the PNP- and KNP-schemes directly, we chose the same boundary condition for the potential in the PNP-scheme.

In order to make the system fully determined, we set the additional requirement

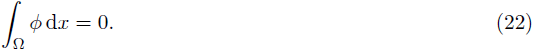

### 4.3 Application details

In the following, we go through the details of the three applications presented.

#### 4.3.1 Application 1: 1-D step concentration

In the first application, which was implemented using the PNP-, KNP-, and DO-schemes, we used an idealized 1-D mesh with a resolution sufficiently fine for PNP to be stable. Two ion species were included in this setup; Na^+^ and X^−^. We present two simulations using this setup.

1. The first simulation was performed on a mesh on the interval [*−*0.1 µm, 0.1 µm]. The mesh had uniformly spaced vertices, with Δ*x* = 0.02 nm. The time step was set to Δ*t* = 0.1 ns. The simulation was stopped at *t*_end_ = 0.1 µs.

2. The second simulation was performed on a mesh on the interval [*−*50 µm, 50 µm], with Δ*x* = 0.01 µm, and *t* = 1.0 ms. The simulation was stopped at Δ*t*_end_ = 10 s.

For both simulations, the initial concentration of both ion species was set to a step function, as

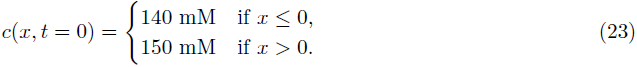

#### 4.3.2 Application 2: Point source/sink simulation

For the second application, which was implemented using the KNP-scheme, we generated a 3-D box-shaped mesh with opposing corners located at (0 µm, 0 µm, 0 µm) and (400 µm, 400 µm and 40 µm). The mesh consisted of 27000 linear tetrahedral cells. A K^+^ point source and a K^+^ point sink were placed in the system. They were placed with a distance of 160 *µ*m apart, at x_1_ = (120 µm, 200 µm, 20 µm) and x_2_ = (280 µm, 200 µm, 20 µm). Both were turned on at *t* = 0 and shut off at *t* = 1 s. The point source/sink pair were given opposites fluxes, of magnitude *±I*(*Fα*)*^−^*1, where *F* is Faraday’s constant, *α* is the ECS fraction, and *I* is the input current, set to *I* = 0.1 nA. The simulation was started at *t*_start_ = 0 s, and stopped at *t*_end_ = 2 s, with a time step of Δ*t* = 2 ms. Two measurement points were chosen to create time series of the potential. These were placed at x_left_ = (120 µm, 205 µm, 20 µm) and xright = (280 µm, 205 µm, 20 µm) (see Results).

#### 4.3.3 Application 3: Input from a morphologically detailed neuron

In the final simulation we modeled the ion sources from a morphologically detailed neuron [48], using the KNP-scheme. The source terms were generated by simulations run on the NEURON simulator. In the NEURON-simulation, the spatial location and magnitude of ion specific membrane currents were stored. This was then used as an external input to the KNP simulation in the form of ion sinks and sources distributed in the 3-D ECS mesh (i.e., ions entering or disappearing from the ECS), as illustrated in Fig 1A-B. The NEURON simulations were run independently from the KNP simulation, which means that there was no feedback from the ECS dynamics to the neurons.

**Fig 1.**
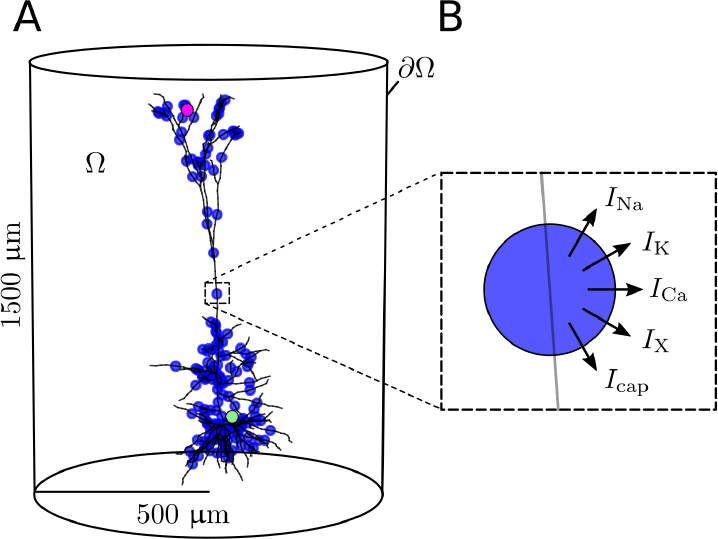
Model system of a 3-D cylinder of ECS containing a morphologically detailed neuron. (**A**) The simulated domain is denoted by Ω, and the boundary is denoted by *∂*Ω. Exchange of ions between the neuron and ECS was modeled as a set of point sources (marked by blue dots). Two measurement points were used to create time series in the Results-section. These are shown as the green and purple points. (**B**) Each point source was included as a sink/source of each ion species, as well as a capacitive current.

In the 3-D case, for a simulated neuron consisting of *N_C_* compartments, with centers at x_1_, x_2_, …, x*_N__C_*, the ionic source terms in the Nernst-Planck equation would be:

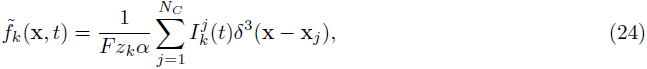

where *δ*^3^(x) is the 3-D Dirac delta function, and 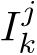(*t*) is the current from compartment *j* specific to ion species *k*, at time *t*. By convention, we define the current to be positive if there is a flow of positive charge out of the cell. In addition to the ionic membrane currents 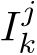 there are also capacitive membrane currents 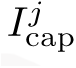. As capacitive sources are not ion specific, they did not appear as source terms in the Nernst-Planck equation (Eq 5), but gave rise to source terms in the KNP equation for the extracellular field. (cf. Eq 10):

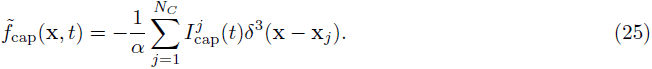

With these source terms (Eqs 24 and 25), the KNP equation (Eq 12) can be written:

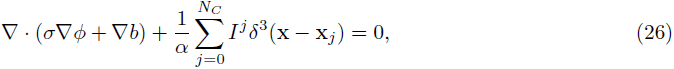

where *I^j^* denotes the sum of all ionic and capacitive membrane current at compartment *j*,

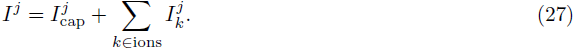

To generate the ion-specific membrane currents, we used a model developed for cortical layer 5 pyramidal cells [48], consisting of 196 sections. All ionic currents, as well as the capacitive current, was recorded for each section. The neuron was driven by Poissonian input trains through 10.000 synapses. The synapses were uniformly distributed over the membrane area, and were tuned to give the model neuron a firing rate of about five action potentials (APs) per second. The NEURON simulation was nearly identical to that used by us previously, and we refer to the original implementation for further details [4]. The only difference is that in the previous paper, the membrane currents were stored in 1-D compartments based on their position in the discretization of the ECS, while in this implementation they were stored as point sources with 3-D coordinates, which allows for greater morphological detail.

The extracellular space was modeled as a 3-D cylinder with a height of roughly 1500 µm and a radius of roughly 500 µm. The mesh was automatically generated using *mshr* [49], yielding a mesh with 53619 linear tetrahedral cells. All ionic concentrations were clamped at the boundary, to their initial background concentrations (cf. Table 2). The time step was Δ*t* = 0.1 s, and the simulation was stopped at *t*_end_ = 80 s. An additional simulation was performed using a smaller time step of Δ*t* = 0.1 ms. Starting with the concentrations found at the end of the 80-second simulation, this secondary simulation was stopped at *t*_end_ = 80.2 s.

Two measurement points were chosen for creating time series of the concentrations. These are referred to as the green and purple measurement points in the Result-section, and are shown in Fig 1A. In the computational mesh, with the soma centered at the origin, the green point was set at x_green_ = (20 µm, 20 µm, 20 µm), and the purple point was set near the apical dendrites, at x_purple_ = (−100 µm, 1100 µm, 0 µm).

## 5 Results

Below we present the results obtained with the three different applications listed in the introduction and methodology. In the first subsection, we have explored the validity of the KNP framework by comparing it to the PNP framework in a simplified application representing a box filled with a solution containing only two ion species, Na^+^ and X^−^. We also showed that even for this system, containing no current sources, a DO model gave inaccurate results. In the second subsection, we have highlighted the differences between the KNP- and VC-schemes using an idealized application containing only a single current source and sink, both mediated by K^+^ ions. Finally, in the third subsection, we have used the KNP-scheme to model the ion dynamics and electric potential around a detailed model of a pyramidal neuron. For this final application, we also compared the predictions of the KNP-scheme to those of the simpler DO- and VC-schemes, and analyzed their differences.

### 5.1 Application 1: Comparison of KNP-, PNP- and DO-schemes for a salt concentration gradient

The differences and similarities between KNP, PNP and DO were illustrated using the simplified 1-D domain representing a box filled with a solution containing two ion species, Na^+^ and X^−^, with an initial concentration gradient (see Methods).

When we simulated Application 1 using the PNP-scheme, two distinct modes of behavior were revealed. Firstly, an initial charge-relaxation mode occurred. During this mode, the X^−^ concentration spread towards the left side of the box faster than the Na^+^ concentration, since X^−^ had a larger diffusion coefficient than Na^+^. Due to the charge separation caused by this process, an electrical potential difference rapidly built up in the system (see blue line in Fig 2A). The charge separation process only went on for a few nanoseconds, before the system reached a quasi-steady state. Secondly, a quasi-steady state occurred, in which the potential difference was so that diffusion and electrical drift were in equilibrium, and further charge separation was prevented. The quasi-steady state potential changed on a slow timescale of seconds (Fig 2B). This was the timescale at which the concentration differences in the system evened out.

**Fig 2.**
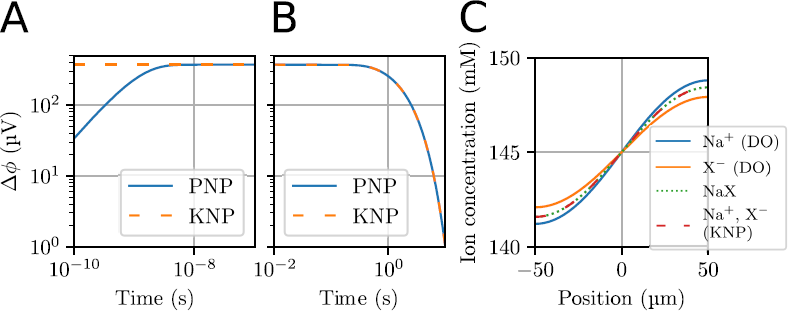
Electrodiffusion in a simple 1-D system containing no current sources. The system had an initial step concentration of NaX being 140 mM for *x <* 0 and 150 mM for *x >* 0. (**A**) Dynamics of the potential difference between the left and right sides of the system on a short timescale. Δ*ϕ* is defined as Δ*ϕ* = *ϕ*(*x*_right_) *− ϕ*(*x*_left_), where *x*_right_ = 0.1 µm and *x*_left_ = *−*0.1 µm. In PNP simulations, the system spent about 10 ns building up the potential difference across the system. In KNP simulations, the steady-state potential was assumed to occur immediately. After 10 ns, the schemes were virtually indistinguishable. (**B**) Dynamics of the potential between *x*_right_ = 50 µm and *x*_left_ = *−*50 µm on a long timescale. The PNP- and KNP-schemes gave identical predictions. (**C**) The concentration profiles in the system at *t* = 1.0 s as obtained with KNP, DO and according to the theoretical approximation when NaX develops as a single concentration moving by diffusion with a modified diffusion coefficient.

A comparison between the two lines in Fig 2A and B highlights the key difference between the PNP (blue line) and KNP (yellow dashed line) schemes. The KNP-scheme does not model the relaxation of the charge concentration, but derives the quasi-steady state potential directly. The results using KNP therefore deviated from those using PNP only on very short timescales during which the quasi-steady state potential was built up. After the initial charge-relaxation (*t >* 5 ns), the two schemes were in excellent agreement.

In theoretical studies of binary systems of ions, it has been shown that the ion concentrations will develop approximately as a single particle species moving by simple diffusion with a modified diffusion coefficient [50],

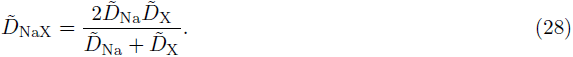

In Fig 2C the ion concentration profiles obtained with this theoretical approximation have been compared to a KNP-based simulation, as well as a DO-simulation where the electric charge of the ions was ignored, so that they were allowed to diffuse independently. The KNP solution was in agreement with Eq 28, and shows that the anion and cation concentrations coincided closely due to the strong electrical forces that would be associated with charge separation. This comparison highlights the shortcomings of the DO-model, which (i) gave inaccurate predictions of the two ion-concentration profiles, and (ii) predicted that the twin ion species were not locally balanced, so that the system contained a charge density *ρ* = *F* (*c*_Na_ *− c*_X_) which varied over the x-axis. According to the Poisson equation (Eq 8), the charge density associated with the concentration profiles in Fig 2C would amount voltage differences of 247 kV across the system, which is clearly not physically realistic.

### 5.2 Application 2: Comparison of KNP- and VC-schemes for an ion source-sink pair

We next compared the KNP-scheme with the simpler VC-scheme for computing ECS potentials. We first did this for an idealized setup, using the computational domain of a 3-D box consisting of the extracellular domain. The initial ion concentrations were uniform, with values from Table 2. The box contained one K^+^ point source, and one K^+^ point sink, with equal magnitudes (see Methods).

Fig 3A shows the electrical potential *ϕ* at the end of the stimulus period (*t* = 1 s). The resulting electrical potential could be split into two contributions: Fig 3B shows the contribution from the membrane currents (the source and sink current), *ϕ*_VC_, while Fig 3C shows the contribution from the diffusive currents in the ECS, *ϕ*_diff_.

**Fig 3.**
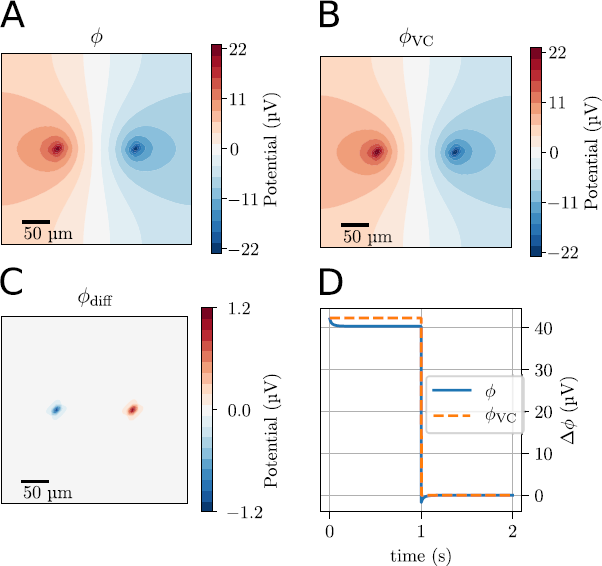
Application with a K^+^ point source/sink pair in a 3-D box containing a standard ECS ion solution. The source and sink were turned on at *t* = 0 s, and turned off at *t* = 1 s. The simulation was stopped at *t* = 2 s. (**A**) The total electrical potential *ϕ* at the end of the stimulus period (*t* = 1.0 s) in a plane intersecting the source and sink. (**B**) The volume conductor component of the electrical potential, *ϕ*_VC_. (**C**) The diffusive component of the electrical potential *ϕ*_diff_. (**D**) The difference in the electric potential between points close to the point sources (blue line), also shown is the differences in the VC component of the potential (stapled yellow line). Δ*ϕ* is defined as Δ*ϕ* = *ϕ*(x_left_) *− ϕ*(xright), where x_left_ and x_right_ were points 5 µm away from the left and right, respectively (see Methods).

In this simulation, *ϕ*_VC_ had a constant value which was reached immediately when the source and sink were turned on, and remained constant throughout the stimulus period since the source and sink were constant. This was not the case for *ϕ*_diff_, which was close to zero at *t* = 0, but increased gradually throughout the simulation, as concentration gradients built up in the system. The diffusion potential *ϕ*_diff_ was quite local, and gave strongest contribution close to the source and sink (Fig 3C).

Application 2 illustrates the limitations with the VC-scheme. Fig 3D shows the potential difference between two points close to the source and sink. *ϕ*_VC_ gave a good estimate of the real potential *ϕ* only for a brief period after the source and sink had been turned on, when the contribution from *ϕ*_diff_ was still negligible. As the simulation progressed, *ϕ*_VC_ became an inaccurate estimate of the true (KNP) potential *ϕ*, since also the diffusive current in the ECS contributed. The diffusion component had a sort of screening effect, reducing the potential difference between the source and sink compared to what we would predict in the absence of diffusion (i.e. *|ϕ| < |ϕ*_VC_*|*. After roughly 0.1 seconds, *ϕ*_diff_ had lowered *ϕ* by about 5 %, compared to what the VC model predicted. This effect lasted until the source and sink were turned off (Fig 3D). After this, *ϕ*_VC_ immediately dropped to zero, and only the effect of *ϕ*_diff_ remained. In the absence of sources, *ϕ*_diff_ decayed gradually until system reached a steady state.

### 5.3 Application 3: Ion dynamics in the ECS of a morphologically detailed neuron

Finally, we used the KNP-scheme to model electrodiffusion in a 3-D cylinder surrounding a morphologically detailed neuron (Fig 1A). The neuron was modeled as a set of point sources and sinks that exchanged ions with the system, as illustrated in Fig 1B (see Methods for details). Since ECS ion concentrations generally vary on a slow timescale (order of seconds), the system was simulated for 80 s. Due to the efficiency of the KNP-scheme, we could run this simulation in less than 15 hours using 4 CPU cores in parallel on a standard desktop computer.

#### 5.3.1 Concentration gradients in the ECS caused by neuronal membrane currents

Fig 4 shows snapshots of the ion-concentration profile surrounding the neuron after *t* = 80.0 s of neural activity. Due to the slow nature of diffusion, the concentration changes clearly reflected the presence of neuronal sources/sinks. The largest changes were seen for the Na^+^ and K^+^ concentrations near the soma and axon hillock. This reflects the strong neuronal output of K^+^ and uptake of Na^+^ from the soma-near regions during action potentials. Animations that show the time evolution of the concentrations were also created (see S1 Animations).

**Fig 4.**
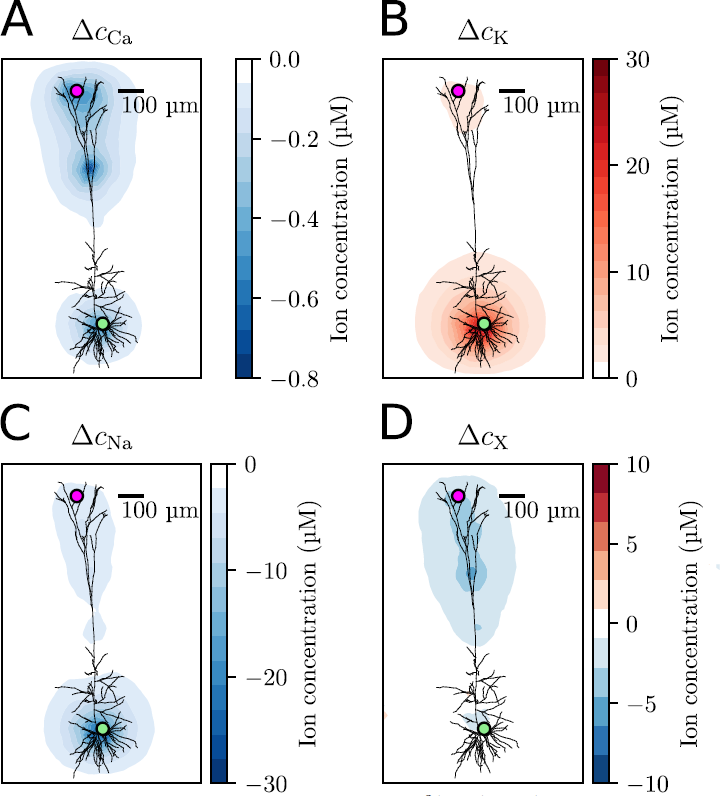
(**A**)-(**D**) The change in ionic concentration for Ca^2+^, K^+^, Na^+^, and X^−^, respectively, as measured at *t* = 80 s, compared to the initial concentration. Δ*c_k_* is the change in concentration from the initial value, defined as Δ*c_k_* = *c_k_*(*t*) *− c_k_*(0). The concentrations were measured in a 2-D slice going through the center of the computational mesh, and the neuron morphology was stenciled in.

#### 5.3.2 The effect of electrical fields on extracellular ion dynamics

In general, ions move both by electrical drift and diffusion. To explore the relative effect of these transport processes on the ECS ion-concentration dynamics, we compared two simulations where (i) ions moved by independent diffusion as predicted by the DO-scheme, and (ii) where ions moved due to electrodiffusion as predicted by the KNP-scheme. The two schemes were compared by looking at time series for the ion concentrations at two selected measurement points, one near the soma (green), and one near the apical dendrites (purple), as indicated in Fig 4. The concentration time series obtained with the KNP-scheme are shown in Fig 5A and B, for the green and purple measurement points, respectively. In the remaining panels of Fig 5, the KNP and DO-schemes are compared.

**Fig 5.**
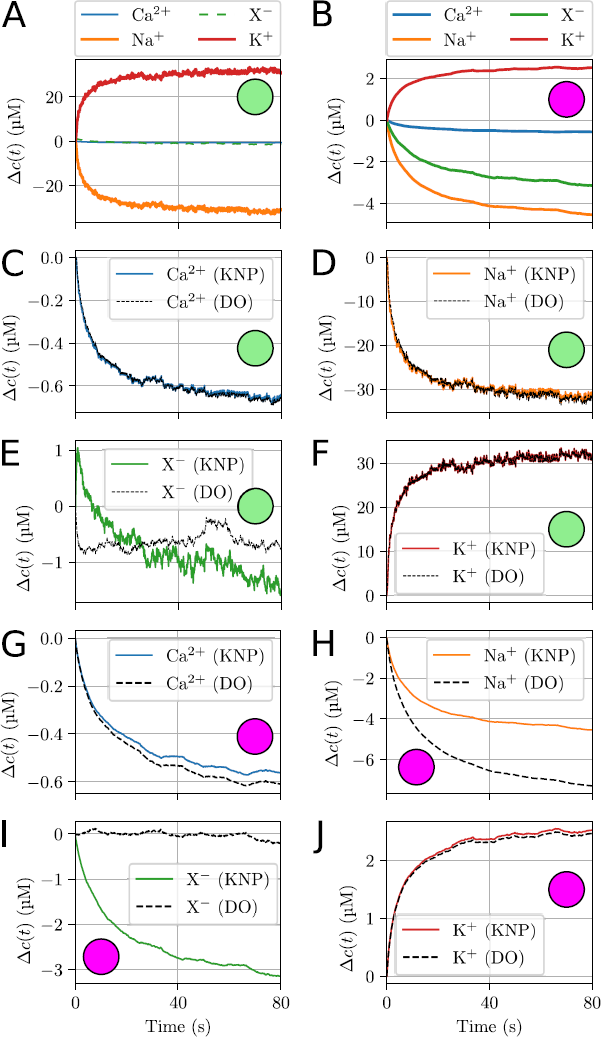
Ion concentration dynamics at selected measurement points. The locations of the measurement points were in the ECS near the soma (green) and apical dendrites (purple) of the neuron. Δ*c*(*t*) is the change in concentration from the initial value, defined as Δ*c*(*t*) = *c*(*t*) − *c*(0). (**A**)-(**B**) Dynamics of all ion species as predicted by the KNP-scheme. (**C**)-(**F**) A comparison between the KNP and DO predictions at the measurement point near the soma. (**G**)-(**J**) A comparison between the KNP and DO predictions at the measurement point near the apical dendrites. (**C**)-(**J**) To generate the DO predictions, we set the ECS field to zero, which ensured that the ECS dynamics was due to diffusion only.

At the measurement point near the soma (Fig 5C-F), the KNP- and DO-schemes gave similar predictions for all ion species except X^−^ (Fig 5E). To explain this, we separate the local concentration dynamics into (i) a component due to neuronal uptake/efflux, (ii) a component due to ECS diffusion, and (iii) a component due to ECS electrical drift. The first two components (i-ii) were shared between the KNP- and DO-schemes, and the differences between the schemes were due to the latter component, which was absent from the DO-scheme. The component (iii) represents ions being electrically forced into a local region to ensure local electroneutrality, and would in principle affect the dynamics of all ion species. The reason why the effect was small in the case of Na^+^ and K^+^, is that the concentration changes of these ions were dominated by (i) the neuronal uptake and output during AP firing, and less so by (ii-iii) electrodiffusion in the ECS. The reason why the relative effect of electrical drift was larger for X^−^ than for Ca^2+^ is twofold. First, X^−^ had a larger diffusion constant than Ca^2+^, and was therefore more mobile (cf. Table 2). Secondly, the number of electrically migrating ions is proportional to the abundance of a given ion species (cf. Eq 4), as well as the valence of the ion species. Even though Ca^2+^ has a larger valence than X^−^, as X^−^ was a significantly more abundant than Ca^2+^ in the ECS (cf. Table 2), we would expect X^−^ to be most affected by electrical drift.

In the apical dendrites, the membrane currents were not dominated by the stereotypical AP exchange of Na^+^ and K^+^, and shifts in the different ion concentrations were more similar in magnitude. At the measurement point near the apical dendrites, the KNP and DO predictions deviated for all ion species (Fig 5G-J). The deviations were largest for Na^+^ and X^−^, which again reflects the fact that these were the two most abundant ion species in the ECS (cf. Table 2).

The exact nature of the ECS dynamics in Fig 5 depended on the specific distribution of ion channels in the selected neuronal model. Although this model choice was somewhat arbitrary, these simulations still serve as a clear demonstration that ECS ion dynamics in general is of electrodiffusive nature, and will be inaccurately modeled by a DO-scheme.

#### 5.3.3 Local effects of diffusion on the extracellular potential

Above, we saw that the ECS potential could influence the local ion-concentration dynamics. Here, we explore how the ion-concentration dynamics can influence the local potential *ϕ*. We first study a low-pass filtered version of *ϕ* by taking the average over a 100 ms interval. Fig 6A shows the spatial profile of the low-pass filtered *ϕ* at *t* = 80 s, while Fig 6B and C show the diffusion component (*ϕ*_diff_) and VC component (*ϕ*_VC_) of *ϕ*, respectively. Evidently, the two components had similar amplitudes, which means that ECS diffusion gave a substantial contribution to the total potential *ϕ*. It should be noted that this conclusion is based on the low-pass filtered version of the signal, and that the higher frequency components are studied below.

**Fig 6.**
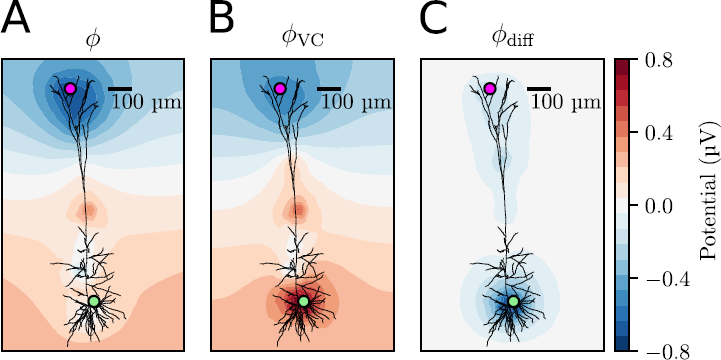
Spatial profile of the ECS potential at a selected time point. The ECS potential was averaged over a 100 ms interval between *t* = 79.9 s and 80 s. This averaging low-pass filters the potential. (**A**) ECS potential as predicted by the KNP-scheme. (**B**) The VC component of the potential. (**C**) The diffusive component of the potential.

Interestingly, the simulations demonstrated that *ϕ*_diff_ was much more local than *ϕ*_VC_. Whereas the membrane currents generated potential fluctuations over the entire simulated region, the *ϕ*_diff_ contributions were confined to the regions of space where concentration gradients were nonzero. Since diffusion is a slow process, only a relatively small region of about 200 µm around the neuron was affected during the 80 s simulation. The local nature of *ϕ*_diff_ implies that *ϕ*_VC_ gives a good estimate of *ϕ* in regions where concentration gradients are small, even if diffusive effects are present in other regions of the system.

To compare *ϕ*_VC_ to other common implementations of the VC-scheme, we note that for other implementations, *σ* is usually kept constant in the entire domain. We measured *σ* throughout the domain for the whole simulation, and found that it varied by about at most 0.001 %, which means that this result is essentially the same as what would be found by the traditional implementation of the VC-scheme, given that the same boundary conditions are used.

#### 5.3.4 Diffusive currents affect the low-frequency part of the LFP

To gain further insight in how diffusion can affect ECS potentials, we also explored the temporal development of *ϕ* at two selected measurement points (marked in green and purple in Fig 6). Fig 7A-B show the dynamics of *ϕ* (blue curve) during an 80 seconds simulation, as well as its components *ϕ*_VC_ (orange curve) and *ϕ*_diff_ (green curve). Again, each data point represents the average over a 100 ms interval, so that *ϕ* was effectively low-pass filtered. As we also saw above, the low-frequency contributions of *ϕ*_VC_ and *ϕ*_diff_ were similar in magnitude (and opposite in sign).

**Fig 7.**
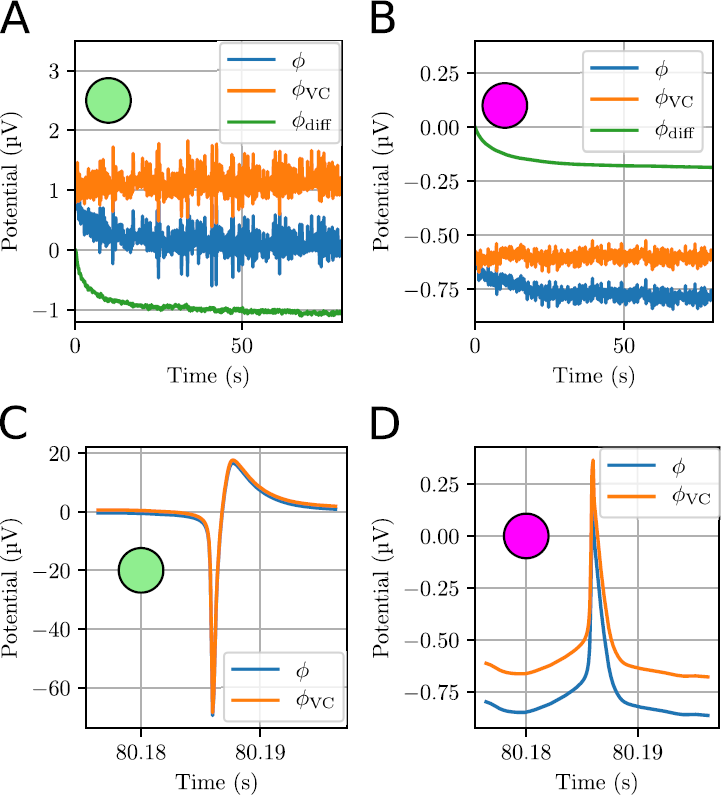
Dynamics of the ECS potential at selected measurement points. The selected measurement points were in the ECS near the soma (green) and apical dendrites (purple) of the neuron. (**A**)-(**B**) Dynamics of the ECS potential for the first 80 s of the simulation. Each data point represent an average over a 100 ms time interval, which effectively low-pass filtered the signal. (**C**)-(**D**) High resolution time series for the ECS potential during a neural AP. The data were not low-pass filtered, but had a temporal resolution of 0.1 ms.

The time series clearly showed that *ϕ*_VC_ varied more rapidly in time than *ϕ*_diff_. Even when low-pass filtered, *ϕ*_VC_ fluctuated several times between about 0.5 and 1.5 µV during the 80 s simulation, while *ϕ*_diff_ underwent an almost monotonous decrease from zero towards -1 µV (Fig 7A).

Due to the slow nature of the diffusive currents, their contribution to the ECS potentials was close to DC-like, and we would therefore not expect diffusion to have an impact on brief signals, such as extracellular AP signatures. We verified this in an additional simulation, which started (with the ECS concentrations) at *t* = 80 s, and which used a smaller time step (0.1 ms) in order to simulate an AP with sufficient temporal resolution. The simulation was run for 200 ms, and an AP was observed at roughly *t* = 80.18 s.

Fig 7C and D shows the time course of *ϕ* (blue curve) and *ϕ*_VC_ (orange curve) during the AP. The constant offset between the two curves was due to the diffusive contribution *ϕ*_diff_, which was accounted for by *ϕ*, but not by *ϕ*_VC_. In the apical dendritic region, the offset was of comparable magnitude to the amplitude of the AP signature (Fig 7C). The diffusion evoked offset was even larger outside the soma, but its relative impact on *ϕ* was smaller, since *ϕ*_VC_ there varied by more than 100 µV during the AP (Fig 7D). The offset was seen to distort the shape of the AP signature at either of these points. Hence, we can conclude that diffusion can have a strong effect on the low-frequency components of *ϕ*, but that the high frequency components are unaffected by diffusion. This conclusion is in line with what we found in previous studies based on simpler, 1-D implementations of the KNP-scheme [4, 20].

## 6 Discussion

In the current work we presented a 3-D version of the electrodiffusive KNP-scheme, and used it to simultaneously simulate the dynamics of ion concentrations and the electrical potential surrounding a morphologically detailed neuron. We demonstrated the validity of this simulation framework by comparing it to the more physically detailed, but more computationally demanding PNP-scheme (Fig 2). Furthermore, we demonstrated the need for an electrodiffusive simulation framework by showing (i) how its predicted ion concentrations deviated from predictions obtained with the diffusion only (DO) scheme, which ignores concentration variations due to electrical drift in the ECS (Figs 2 and 5), and (ii) how its predicted potentials deviated from predictions obtained with the volume conductor (VC) schemes, which ignores voltage variations due to diffusive currents in the ECS (Figs 3, 6, and 7).

To our knowledge, the presented model is the first in the field of computational neuroscience that can handle electrodiffusive processes in 3-D on spatiotemporal scales spanning over millimeters and seconds without demanding an insurmountable amount of computer power. Even the most resource-demanding simulations presented here could be performed in about 15 hours on a normal stationary computer, and we believe that simulation efficiency can be improved even further if we select an optimal numerical scheme for KNP. In the current work, all simulations were run using an implicit Euler time-stepping scheme. This choice was mainly based on the requirements of the PNP-scheme, which requires an implicit scheme in order to not become unstable. The KNP-scheme is, however, much more stable than the PNP-scheme, and in future KNP implementations we will investigate the possibility of using an operator-splitting approach for the numerical solver, which would likely improve the efficiency of the simulations.

### 6.1 Model limitations

The presented implementation of the 3-D KNP-scheme was limited to a confined ECS region surrounding a single pyramidal neuron modeled with the NEURON simulator. Although this approach provided clarity regarding the possible single-neuron contribution to the ECS electro- and concentration dynamics, there are many reasons why this simulation setup was far from representative for any biologically realistic scenario.

Firstly, the brain is packed with neurons and glial cells, and even the small volume considered here would in reality be populated by hundreds of neurons that would contribute to the ECS dynamics. In a realistic scenario, ECS concentration shifts may become much larger than for the single-neuron case, where the maximal changes in ion concentration were roughly 1 % (for K^+^ near the soma).

Secondly, the simulations were performed under the simplifying assumption that there was no feedback from the ECS dynamics to the neurodynamics. In reality, shifts in extracellular (or intracellular) concentrations would influence the reversal potential of transmembrane currents, which would have an impact on the neurodynamics (see [51]). In addition, simulations based on the NEURON simulator do not account for ephaptic feedback from extracellular fields onto neurons, which could also have moderate impact on neurodynamics (see [52]).

Thirdly, the neuron model used in our study did not include the Na^+^/K^+^-exchanger pump [48]. This is the case for most neuron models currently available in NEURON (for a model with ion pumps, see [53]). Together with astrocytic uptake mechanisms, the exchanger pump would strive towards maintaining extra- and intracellular ion concentrations close to baseline levels. In the presence of such mechanisms, the single-neuron contribution to ECS concentration shifts would likely be smaller than in the simulations presented here, or would require a higher neural activity level in order to occur. The model limitations mentioned above were also present in the previous 1-D implementation of the KNP-scheme, and we refer to this previous work for a more thorough discussion [4].

### 6.2 Previous models of ECS electrodiffusion

Several previous studies have explored ECS electrodiffusion on small spatiotemporal scales [19, 24–30, 32–35]. Electrodiffusive models tailored to explore larger spatiotemporal scales have to our knowledge so far only been implemented in 1-D [4, 20, 36–38].

We have previously used a 1-D implementation of the KNP-scheme to explore the effect of diffusive currents on ECS potentials [4]. Qualitatively, the results from the 1-D simulations were similar to those found in the current 3-D model, and in both cases it was concluded that diffusive currents affected the low-frequency components of ECS potentials (Fig 7). In the 1-D implementation, diffusion and electrical drift were confined to the spatial direction along the neuronal extension, i.e. along the basal dendrite-soma-apical dendrite axis, and did not occur in the lateral direction. As discussed in [4], the 1-D assumption is equivalent to assuming lateral homogeneity, which means that any neuron is surrounded by neighboring neurons with identical activity patterns as itself. Clearly, this is would not apply to most biological scenarios, which means that the results obtained with the 3-D implementation are generally more reliable. In contrast to the 1-D model, the 3-D model predicts how ion concentrations vary in the direction laterally from the neuronal sources, and can therefore be applied to new biological problems such as spreading depression, which is associated with massive redistribution of ions that spreads laterally in the cortex [3, 7, 54].

The diffusion potentials addressed by the KNP-scheme are those that arise due to ECS concentration gradients on relatively large spatial scales. Diffusion potentials of this kind are often referred to as liquid junction potentials, since they are most pronounced at the boundary between two solutions of different ion composition [31, 55]. These diffusion potentials are unrelated to filtering effects hypothesized to arise due to diffusion in the vicinity of the membrane when electric charge is transferred from the intracellular to the extracellular space [56, 57].

#### 6.3 Outlook

The presented version of the KNP-scheme was developed for use in a hybrid simulation setup where the dynamics of ion concentrations and the electrical potential in the ECS were computed with KNP, while the neurodynamics was computed with the NEURON simulator tool. By necessity, this scheme shares the limitations of the NEURON simulator in terms of handling intracellular ion dynamics, which by default is not electrodiffusive in the NEURON environment [10]. A natural future endeavor will therefore be to derive a computational scheme that in a consistent way couples both the intra- and extracellular ion-concentration and voltage dynamics based on the KNP-scheme. Such a scheme will represent a generalization of the previously developed *extracellular-membrane-intracellular* (EMI) model [52, 58], which in a consistent way couples the intra- and extracellular electrodynamics, but so far does not include ion-concentration dynamics and thus not diffusive currents.

This being said, the hybrid KNP/NEURON version presented here is valuable in its own right, since it allows the KNP framework to be combined with the many models that are already available in the NEURON software. Although we here only studied a single neuron’s contribution to the ECS electro-and concentration dynamics, we envision future applications of the 3-D KNP to more complex systems. The 3-D version of the KNP-scheme is generally applicable, and could be used to simulate the dynamics of ion concentrations and the electrical potential in the ECS surrounding arbitrary complex models of neurons or populations of neurons. It could, for example, be used to account for extracellular transport processes in the Blue Brain Simulator [59]. If combined with a cellular system of neurons and glial cells tailored to represent a specific experimental condition, the KNP-scheme could be used to explore the mechanisms behind spreading depression [3, 7, 54], epilepsy [5, 60], or other pathological conditions associated with large shifts in extracellular ion concentrations.

## Supporting information

S1 Animations. Animations of the ion-concentration dynamics. Animations of the temporal evolution of the ion concentrations were created using ParaView. The neuron morphology was stenciled in.

